# The HuBMAP Framework for Advancing Data FAIRness

**DOI:** 10.64898/2026.06.01.728946

**Authors:** Stephen A. Fisher, Josef Hardi, Richard Morgan, Erik Nordgren, Peter M. Kant, Brendan Honick, Jean Rosario, Martin J. O’Connor, Philip D. Blood, DCWG Members, Jonathan C. Silverstein, Mark A. Musen

## Abstract

Since publication of the FAIR Guiding Principles in 2016, the scientific community has increasingly sought to make experimental data findable, accessible, interoperable, and reusable. Operationalizing the FAIR principles in routine scientific workflows remains challenging without a standardized, workable infrastructure. With over 10,000 datasets from over 40 institutions, spanning more than 50 diverse assay types ranging from single-cell sequencing technologies to 2D and 3D spatial omics, the U.S. National Institutes of Health (NIH) Human Bio-Molecular Atlas Program (HuBMAP) consortium has been ideally situated to create a FAIR ecosystem. With the goal of achieving data “FAIRness,” HuBMAP developed and implemented well-defined, community-endorsed metadata reporting standards across the research lifecycle. These reporting standards include detailed schemas, harmonized across a multitude of assays, that define the metadata associated with a dataset and the organization of the corresponding data files. These standards ensure documentation of the data collection process, of the data themselves, and of the manner in which the data are packaged for sharing, while remaining compliant with the Health Insurance Portability and Accountability Act (HIPAA). The use of these reporting standards, in tandem with technology to foster adherence, allows HuBMAP to fulfill its goal of generating FAIR data for open dissemination through its Data Portal and Human Reference Atlas. The procedures and simple workflow adopted by HuBMAP investigators serve as a model for other scientific communities aiming to maximize the value of varied datasets addressing a shared research question. The HuBMAP end-to-end, metadata-centered workflow has been replicated and enhanced by the NIH Cellular Senescence Network (SenNet) consortium and is readily available through open-source technology for others to utilize.

## 1. Introduction

Within the scientific community, there is broad recognition of the importance of making research data findable, accessible, interoperable, and reusable (FAIR). Further, new and expanding governmental mandates require investigators not only to share their datasets but also to ensure that the datasets are FAIR. The value proposition set out by the FAIR Guiding Principles published in 2016 has been partially realized through important secondary analyses of other people’s FAIR data, significantly increasing their value. Yet despite the strong appetite and requirements for generating FAIR data, confusion and operational challenges around data collection and dissemination have stymied the realization of potential benefits^1^.

A common misperception exacerbates this problem: investigators can declare that their data are FAIR because they have deposited the data in online repositories^2^. Unfortunately, these data are often accompanied by only the minimal metadata needed to obtain a digital object identifier (DOI) for their contribution. More often than not, such metadata are insufficient to support reproducibility because they either lack essential information or fail to conform to standardized data-annotation schemas. In contrast, when metadata are comprehensive and accurate (“rich”), with a greater number of searchable metadata fields, following a pre-determined structure, the datasets become inherently more findable, and it is easier to elucidate experimental conditions, rendering the data more interpretable, interoperable, and reusable^3^.

Metadata, or “data about data,” are commonly divided into at least three distinct categories: (1) descriptive metadata, (2) structural metadata, and (3) administrative metadata^4^. Descriptive metadata describe the source of the data—in our case, this includes details about the type of experiment that was performed, the experimental conditions, and the subject of the experiment. Structural metadata convey how the data are organized—how the datasets are packaged, what files are included, and how the data are encoded. Administrative metadata include information needed to manage the data, such as creation dates and licensing restrictions. Descriptive metadata, which include standardized attributes linked, where appropriate, to controlled value sets and ontologies, are essential for FAIR data because they enable third parties to understand how datasets were generated and how they should be interpreted. Descriptive metadata must conform with such standards for datasets to be *findable* and *interoperable* with related datasets. Structural metadata are required to understand how datasets are assembled physically, enabling *accessibility*. Taken together, descriptive and structural metadata ensure that datasets are *reusable*.

In this paper, we report how a consortium of life-science investigators has developed internal standards for descriptive and structural metadata, and general workflows for authoring, validating, and applying those standards to ensure the FAIRness of its datasets. The approach builds on trends that have emerged over the past 30 years in the biomedical community that emphasize the use of community-based, descriptive metadata standards to support data sharing and reuse. Several groups have come together to create well-known standards, such as the Minimum Information About a Microarray Experiment; MIAME^5^. MIAME established a community *reporting guideline* that standardized the descriptive metadata required to interpret, verify, and share microarray experiments. Although the FAIRSharing resource^6,7^ reports more than 300 such community-based reporting guidelines, most areas of science and most experiment types still lack standards for descriptive metadata, making data sharing and reuse difficult.

The U.S. National Institutes of Health (NIH) Human Bio-Molecular Atlas Program (HuBMAP)^8–11^ is an expansive research endeavor aiming to identify biomarkers that characterize every type of cell in the healthy human body and to create a comprehensive, three-dimensional mapping of cells to the genes, proteins, lipids, and other substances that identify them. Achieving the goals of HuBMAP requires the accession and integration of thousands of diverse datasets, from over 40 affiliated NIH awardees, studying over 30 tissues and their associated cell types, using more than 50 distinct assay methods and technologies. This massive undertaking is possible only because the data describing cells and their biomarkers are FAIR, allowing the numerous HuBMAP datasets to be stored in a common repository that can be integrated, queried, and presented to users through an interactive portal with a uniform user interface. As this paper describes in detail, HuBMAP translates the FAIR principles into enforceable, machine-actionable community standards that govern documentation of data provenance, experimental conditions, and file organization across the full experimental lifecycle.

## 2. The Working Group model

The HuBMAP Consortium Data Coordination Working Group (DCWG), comprising data providers, data curators, bioinformaticians, and software engineers, is responsible for defining and maintaining standards for data ingestion, storage, access, and visualization. These standards, which support the HuBMAP aim of providing FAIR data, are developed through a collaborative and iterative process involving subject matter experts (SME), ontology specialists, and computational scientists (**Fig. 1**).

**Figure 1.**
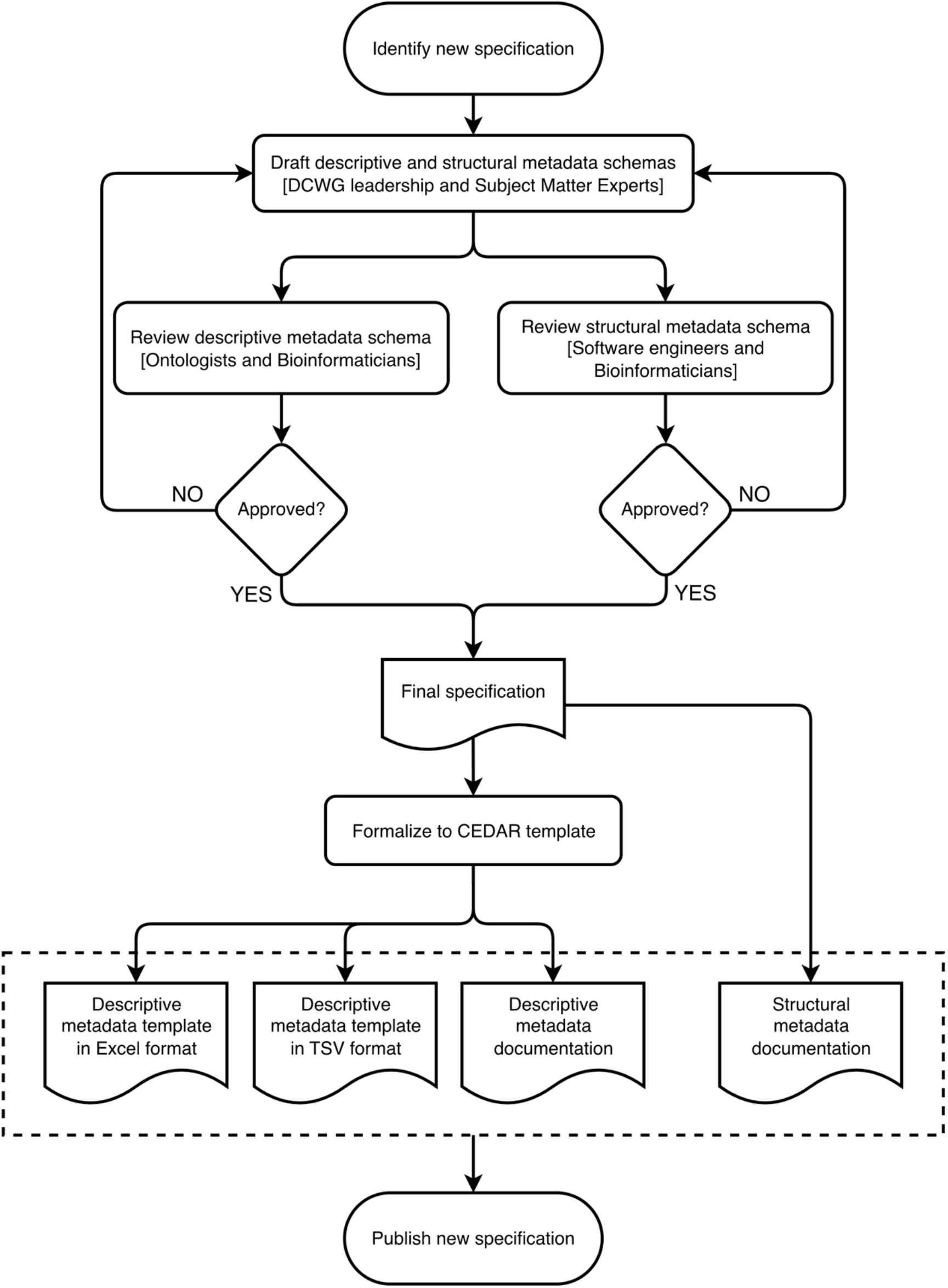
Workflow for development and publication of HuBMAP data standards. New specifications are initiated by the Data Coordination Working Group (DCWG) and subject matter experts, followed by iterative review of descriptive and structural metadata schemas by ontology specialists, software engineers, and bioinformaticians. Finalized specifications are implemented in the CEDAR Workbench and released as standardized templates and documentation for consortium-wide use.

New specifications are initiated in response to emerging assay types or unmet data requirements. The DCWG leadership works with SMEs to draft both assay-specific schemas for descriptive metadata and structural metadata specifications, which together define required and optional metadata attributes, naming conventions, and file organization. Ontology experts and bioinformaticians then review the descriptive metadata schemas and map the datatypes of metadata attributes to established ontologies and value sets, introducing new terms where necessary. In parallel, software engineers and bioinformaticians evaluate the structural file specifications (against SME-provided representative datasets) to ensure that they support downstream processing, visualization, and integration. These steps are typically iterative, with feedback at each stage informing successive refinements.

Once finalized, descriptive metadata specifications are implemented as templates in the CEDAR Workbench (see **Section 4**) and documentation is generated for the new descriptive and structural metadata, which are then released for consortium-wide use. This process enables data providers to generate, validate, and submit datasets that rigorously conform to standardized representations.

The DCWG maintains and updates specifications over time, distinguishing between minor and major revisions. Minor updates, such as the addition of optional fields or extensions to controlled vocabularies, are approved and released by DCWG leadership. Major updates, including adding or removing required metadata fields, are developed in consultation with SMEs and require broader review prior to adoption. Data curators play a key role in identifying the need for updates, particularly when similar assays are implemented differently across data provider sites.

## 3. The HuBMAP Metadata Reporting Standards

The HuBMAP program developed community-based metadata reporting standards to operationalize the FAIR principles across thousands of datasets spanning dozens of diverse assay types. HuBMAP also established mechanisms to assist investigators in using these standards in their routine data workflows, ensuring that all datasets adhere to the standards. HuBMAP adopts a comprehensive yet straightforward strategy that employs community-defined reporting standards across the data lifecycle — from sample collection and data generation to dissemination. We propose that other investigators also apply these standards, either by directly adopting them when possible, or by implementing new reporting guidelines, when necessary, with the support of open-source tools.

The HuBMAP metadata standards comprise three integrated components:

1. **Provenance model**. A graph-based model describes the relationships between each stage of an experiment, from donor selection to specimen preparation to assay results, enabling clear tracking and documentation of samples and datasets across the experimental workflow.
2. **Descriptive metadata schemas**. Each stage in the provenance model includes a comprehensive schema that defines its required types of descriptive metadata, providing a standardized description of samples, experiments, and processing context.
3. **Structural metadata schemas**. Standardized hierarchical file and directory specifications, including file type and naming conventions. These specifications ensure that datasets from all providers are organized or packaged consistently, and thus may be readily accessed, understood, and reused by data consumers.

### 3.1 Provenance model: The overarching framework for descriptive metadata

Many scientific datasets arise from a sequence of sample-processing and assay steps that define the context in which measurements are made. The HuBMAP project defines six distinct stages in its particular data collection workflows — (1) identification of a human donor, (2) selection of an organ to study, (3) creation of a tissue block, (4) creation of a tissue section, (5) creation of a tissue suspension, and (6) performing an assay — each step representing different levels of biological material processing (**Fig. 2**). While we elucidated six steps in HuBMAP, other areas of science may require more or fewer steps. These six stages and their relationships (modeled as a graph structure) underlie the workflow captured in the provenance model. This model provides the framework for the descriptive metadata schemas, which capture the types of metadata required for the data that are specific to each stage in the provenance model.

**Figure 2.**
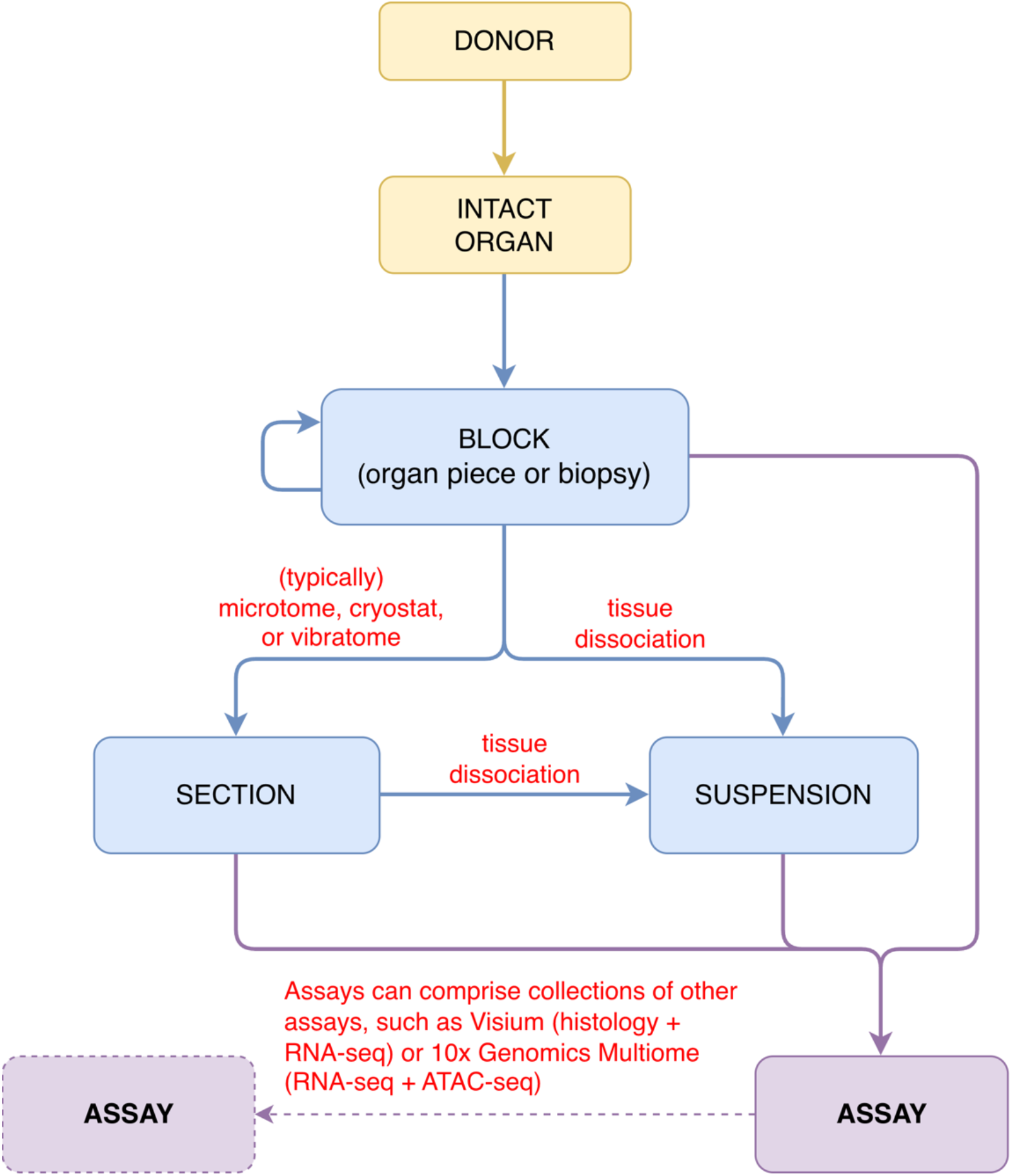
HuBMAP provenance model, represented by a directed-graph structure. The provenance model includes a node for each stage in the data-collection workflow, from donor sampling, to tissue preparation, to data acquisition from one or more assays. Each arrow represents one or more provenance events (e.g., a block may be divided into multiple sections, and a given tissue sample may be subject to more than one assay).

In the provenance model, three of the steps refer to types of “samples”, namely: tissue blocks, tissue sections, and cell suspensions (**Supplementary Description 1**). Tissue blocks originate as one or more pieces of a dissected organ or a biopsy. Blocks may be subdivided into smaller blocks, they may be sectioned multiple times, and they may be converted into suspensions. We do not model biological sources such as cell cultures or organoids, but it would be straightforward to expand the provenance model to include these entities, if required. For example, an organoid could be used in place of an “intact organ”. Similarly, a cell culture can be derived from an organ, organoid, or block. The cell culture could in turn be dissociated into a suspension or processed into a block. This characterization provides a framework for the capture of cell-culture–specific or organoid-specific metadata.

An assay can be applied to tissue blocks (e.g., light sheet microscopy), sections (e.g., 2D imaging), or suspensions (e.g., sequencing), while a single application of an assay can comprise multiple such units (e.g. multiple sections on a single spatial slide). At the assay level, we identified 52 different assay types, and we treat each as a unique node in the provenance model. In the model, assays may be composed of other assays. For example, a sequencing assay might contain both RNA and DNA components, each of which could be run as an independent assay, or *in situ* multi-omic assays could comprise both RNA and protein measurements in each cell.

Beyond traceability and association (the graph enables queries such as finding all datasets from a given donor), the provenance model enables programmatic routing of datasets through assay-specific analysis pipelines. Because each node in the model represents a typed biological entity and assay context, the metadata attached to that node are used to automatically determine the appropriate downstream processing workflow and parameterization (**Supplementary Table 1**). For example, HuBMAP has three different processing workflows for single nucleus ATACseq assays, and the values for the “barcode size” and “barcode offset” determine which of the three processing workflows is used. In this way, provenance is not merely descriptive, but computationally actionable.

### 3.2 Descriptive metadata schemas: Ontology-backed models for humans and computers

Well-defined, “rich” metadata specifications are essential for making datasets FAIR, justifying investment of significant effort in the development of such specifications. With an extensive provenance model and associated metadata-management processes and technologies in place, building metadata schemas (**Table 1**) is a straightforward but arduous community effort across the HuBMAP Consortium—from data providers, to data handlers, to data processors, and, finally, to data-visualization experts. One design goal of the HuBMAP metadata schemas is to eliminate ambiguity among closely related assay variants, particularly in high-throughput sequencing domains where subtle protocol differences materially affect downstream computational processing (**Supplementary Table 2**). The schemas are therefore constructed to describe experiments in sufficient detail to unambiguously determine the correct analysis pipeline and parameter set whenever possible. In particular, assay metadata include details that (1) might vary across datasets, (2) might be used to distinguish among assay variants, and (3) might indicate which analysis pipeline to use to process a given dataset.

**Table 1.** An abbreviated example metadata schema, showing a subset of all fields included in the metadata schema for RNA sequencing. Refer to **Supplementary Table 2** for the full RNA-seq schema.

| Field Name | Description |
| --- | --- |
| parent sample id | The unique identifier from HuBMAP or SenNet for the sample (such as a block, section, or suspension) used to perform the assay. For instance, in an RNAseq assay, the parent sample would be the suspension, while in imaging assays, it would be the tissue section. If the assay is derived from multiple parent samples, this field should contain a comma-separated list of identifiers. Example: HBM386.ZGKG.235, HBM672.MKPK.442 |
| preparation protocol doi | The DOI for the protocols.io page that details the assay or the procedures used for sample procurement and preparation. For example, in the case of an imaging assay, the protocol may start with tissue section staining and end with the generation of an OME-TIFF file. The documented protocol should also include any image processing steps involved in producing the final OME-TIFF. Example: https://dx.doi.org/10.17504/protocols.io.eq2lyno9qvx9/v1 |
| acquisition instrument vendor | The company that manufactures or supplies the acquisition instrument. An acquisition instrument is a device equipped with signal detection hardware and signal processing software. It captures signals produced by assays, such as variations in light intensity or color, or signals corresponding to molecular mass. If the instrument was custom-built or developed internally, enter "In-House". Example: Illumina |
| acquisition instrument model | The specific model of the acquisition instrument, as manufacturers often offer various versions with differing features or sensitivities. These differences may be relevant to the processing or interpretation of the data. If the instrument was custom-built or developed internally, enter "In-House". If the model is unknown, enter "Unknown". Example: NovaSeq X |
| source storage duration value | The length of time the sample was stored prior to processing it. For assays performed on tissue sections, this refers to how long the tissue section (e.g., slide) was stored before the assay began (e.g., imaging). For assays performed on suspensions, such as sequencing, it refers to how long the suspension was stored before library construction started. Example: 12 |
| source storage duration unit | The unit of measurement used to specify the source storage duration value. Example: hour |
| barcode offset | The position within a sequencing read where a barcode sequence begins. In sequencing libraries, barcodes—short DNA sequences used to tag individual cells or molecules—are often arranged in a pattern where each barcode is separated by a constant sequence of known length. These constant sequences create predictable spacing between barcodes, known as offsets. For example, if barcodes are 8 base pairs (bp) long and separated by 30 bp constant regions, the barcodes might start at positions 0, 38, and 76 within the read. If the source material is not barcoded, this field should be marked as "Not applicable". Example: 0,38,76 |
| barcode read | The sequencing read file contains the cell or capture spot barcode sequences. This information is essential when constructing sequencing libraries with a non-commercial kit, as it ensures that barcodes are correctly extracted during data processing. If the source material is not barcoded, this field should be marked as "Not applicable". Example: Read 1 (R1) |
| barcode size | The length of each cell or capture spot barcode in base pairs (bp). If the source material is not barcoded, enter "Not applicable". Example: 16 |
| library layout | Indicates whether the sequencing library was prepared for single-end or paired-end sequencing. This information determines how reads are generated and aligned during data processing. Example: single-end |
| library preparation kit | The reagent kit used to construct the sequencing library, including both the kit name and, if available, version and product number. If the library was prepared using a custom protocol, enter "Custom". Example: 10X Genomics; Chromium Next GEM Single Cell 5' Kit v2, 16 rxns; PN 1000263 |

Each metadata schema comprises descriptions of a rich set of attribute–value pairs, which create a reporting guideline enumerating the standard characteristics of the assay, the specimen, and the experimental context. The permitted values for each attribute have a defined datatype, which may be a simple type such as integer or Boolean, or may denote a nominal entry from the HuBMAP Research Attributes Value Set (HRAVS)^12^. The HRAVS is a custom, project-specific value set that we built by manually curating terms from established biomedical ontologies and vocabularies, including the NCI Thesaurus (NCIT), the Unit Ontology (UO), the Ontology for Biomedical Investigations (OBI), Chemical Entities of Biological Interest (ChEBI), and Logical Observation Identifiers Names and Codes (LOINC), as well as broader biomedical resources such as Medical Subject Headings (MeSH) and Research Resource Identifiers (RRIDs). HRAVS consolidates the categorical fields and their permissible values used across all HuBMAP metadata schemas. For cases where a needed concept has no suitable counterpart in any of these sources, we mint a new HRAVS code for the concept label. In all cases, modeling and ontology specialists review the schema field names, descriptions, and proposed value sets for consistency across all HuBMAP metadata schemas. Since the metadata schemas are extensive, we adopted and expanded the capabilities of the CEDAR Workbench^13^ to ease the burden of completing and validating all required metadata (see **Section 4**).

HuBMAP typically captures donor-level metadata from electronic health record (EHR) systems and extracts the metadata as required and available. For sample-related metadata, there are separate metadata schemas for each of the three sample types (block, section, and suspension), as dictated by the provenance model (**Supplementary Table 3**). When designing metadata schemas, we harmonize fields across schemas as much as possible. For example, all three sample types share some common fields, such as “preparation medium” and “source storage time”. There are also fields that are specific to certain sample types. For instance, the “area” field is only relevant to tissue sections, whereas the “is suspension enriched” field applies solely to suspensions.

We created detailed metadata schemas for each assay type. In addition to identifying metadata common to all assay types, these schemas include fields specific to individual assay classes. For example, “preparation protocol doi” captures the DOI for the assay protocol, as published in https://protocols.io^14,15^, which is relevant to all assays; whereas “slide id” captures a unique ID label for each slide used, and is specific to imaging assays. Fields are harmonized across schemas wherever possible, supporting consistency in data description while preserving assay-specific detail. For imaging-based assays, HuBMAP metadata schemas are further harmonized with organ mapping antibody panels (OMAPs), which define validated antibody panels for tissue mapping^16^. This alignment supports antibody validation, improves reproducibility, and facilitates cross-study comparability in spatial imaging experiments. This approach supports harmonization efforts and simplifies ontology mappings. Comprehensive documentation of the fields and allowed values for all metadata schemas is provided in **Supplementary Table 3**.

Some metadata serves to link entities in the provenance model, such as capturing the temporal relationship between entities. This relational information is important, since, for example, the length of time that a tissue suspension remains frozen before sequencing, or that a tissue block is stored prior to sectioning, may affect downstream analysis results. With human subjects, metadata tracking *dates* (such as when an organ was dissected, or when a sample was sequenced) can be interpreted as not meeting the Health Insurance Portability and Accountability Act (HIPAA) Safe Harbor standard^17^. We therefore designed the metadata schemas to capture the *durations* of sample processing and the intervals between workflow stages, eliminating the need to record any dates. In order to encode the workflow timeline (**Fig. 3**), we capture the temporal duration between each node in the provenance model (i.e., “source storage duration”) and how long a sample was processed within each node (i.e., “processing time”). Although recording time intervals instead of specific dates limits the ability to associate datasets with laboratory-specific events that may affect interpretation or reproducibility (such as personnel, reagent or process changes), this design preserves donor privacy by complying with the HIPAA Safe Harbor requirements mandated by Consortium policy. See **Supplementary Description 2** for examples of temporal provenance.

**Figure 3.**
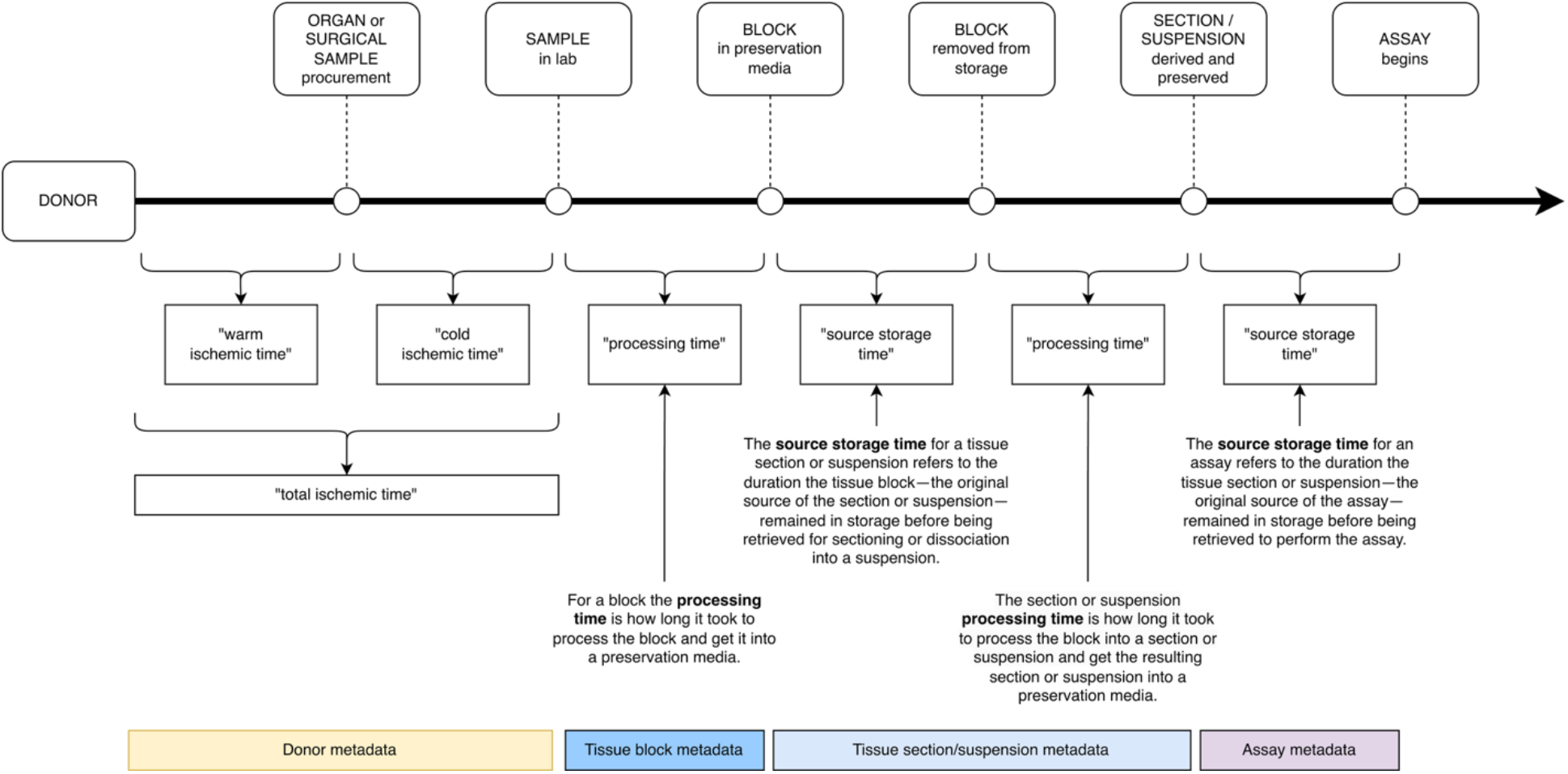
Workflow timeline. Metadata schemas capture how long it takes to process tissue samples within each stage of the provenance model (i.e., “processing time”), as well as how long the tissue samples are stored between the stages (i.e., “source storage time”). The processing and storage durations are typically captured as minutes, hours, or days.

The sequencing domain in particular has many assays that involve similar technologies or that have similar characteristics, and care must be taken to avoid ambiguous metadata, since small experimental differences can affect downstream pipeline processing. For example, when sequencing RNA, one assay might pull RNA from the nucleus and cytoplasm while another might be selective for cytoplasmic RNA. Some RNA sequencing technologies uniquely label the RNA from each cell, whereas other RNA sequencing technologies are not that specific. Furthermore, when preparing the RNA for sequencing, some technologies create duplication artifacts, necessitating further processing to make corrections in the analysis stage. When working with RNA sequencing metadata, the specifications are able to differentiate between current, commonly used assay technologies (**Supplementary Table 1**). This distinction is further relied upon when determining how to process each RNA sequencing dataset. For example, “assay input entity” denotes whether the data are derived from single-cell or single-nucleus entities. At the same time, RNA sequencing technologies that use unique molecular identifiers (UMI) will include “umi read” and “umi size” field values to denote the location and size of the UMIs. Well-specified descriptive metadata can uniquely match datasets with processing pipelines and the parameters required for those pipelines. This nuanced approach underscores the importance of meticulous metadata design in advancing scientific inquiry.

### 3.3 Structural metadata schemas: Standardized file organization

Schemas for descriptive metadata that adhere to standard reporting guidelines and that are linked to standard ontologies are essential for data to be FAIR, but they are insufficient. When data are shared or archived, they need to be packaged using file and directory structures that are systematic and well documented. Placing files with unknown naming conventions and data formats into arbitrary folders or buried in a series of directories is problematic for groups wishing to reuse the data. Thus, in addition to standardized descriptive metadata schemas, ontologies, and value sets, the HuBMAP metadata reporting standards include structural guidelines for the organization and naming of the data files and directories in the corresponding dataset. File and directory specifications establish a harmonized, hierarchical structure to organize and share data files and their associated descriptive metadata in a systematic fashion, which enables automated validation and consistent downstream consumption. Beyond facilitating conventional data reuse, these structural metadata make HuBMAP datasets readily accessible to automated tools and emerging AI-driven systems, which benefit from consistent, machine-interpretable organization of both data and metadata.

Where possible, harmonized file and directory specifications are created and used across similar data types. For example, most assays that generate a 2D or 3D image use the same OME-TIFF^18–20^ image file type with standardized metadata in the OME-TIFF file XML header (**Supplementary Table 4**), placed in a consistent location in the dataset directory tree, and with the same file naming convention, irrespective of the imaging assay used to generate the image. Specifications denote optional and required files and directories, as well as file formats. Files and directories are represented by regular expressions denoting the expected name or naming convention and location (**Table 2** and **Fig. 4A**). Similarly, file formats are either specified as industry-accepted file types (e.g., FASTQ) or manually described (for examples, see **Supplementary Table 5**). This specification allows for automated structural validation of datasets, for example, confirming the existence of all required files in the correct locations, or ensuring that a required data file has the expected format (e.g., validating that a specific CSV file contains the appropriate fields). While data providers are welcome to include additional files and directories, these structural specifications guarantee a minimum set of key files expected for each assay type, in a consistent, harmonized compilation. In this way, data consumers can easily understand and process HuBMAP data files, regardless of which data provider produced and submitted the data.

**Table 2.** File organization schema for the RNA-seq assay. Each assay type has specifications for the hierarchical file structure, file types and naming conventions, with elements shared across dataset types when feasible (e.g., unprocessed data are always situated in the “raw/” directory). In this example, FASTQ files (a common format for storing RNA-seq data) are required and must reside in the “raw/fastq/RNA/” directory. An optional “expected cell count.txt” file can also be included in the “extras/” directory.

| File and Directory Format | Required | Description |
| --- | --- | --- |
| extras/expected_cell_count.txt | No | The expected cell counts for the RNA sequencing dataset. This is an optional file that, if present, will be used by the HIVE's RNA sequencing analysis pipeline. With some datasets, knowing the expected cell count has improved the output of the HIVE analysis pipeline. |
| raw/ | Yes | All raw data files for the experiment. |
| raw/fastq/ | Yes | Raw sequencing files for the experiment. |
| raw/fastq/RNA/ | Yes | Directory containing FASTQ files pertaining to RNA-seq sequencing. |
| raw/fastq/RNA/*R{1,2}*.fastq.gz | Yes | This is a GZip'd version of the forward and reverse FASTQ files from RNA-seq sequencing (R1 and R2). |

**Figure 4.**
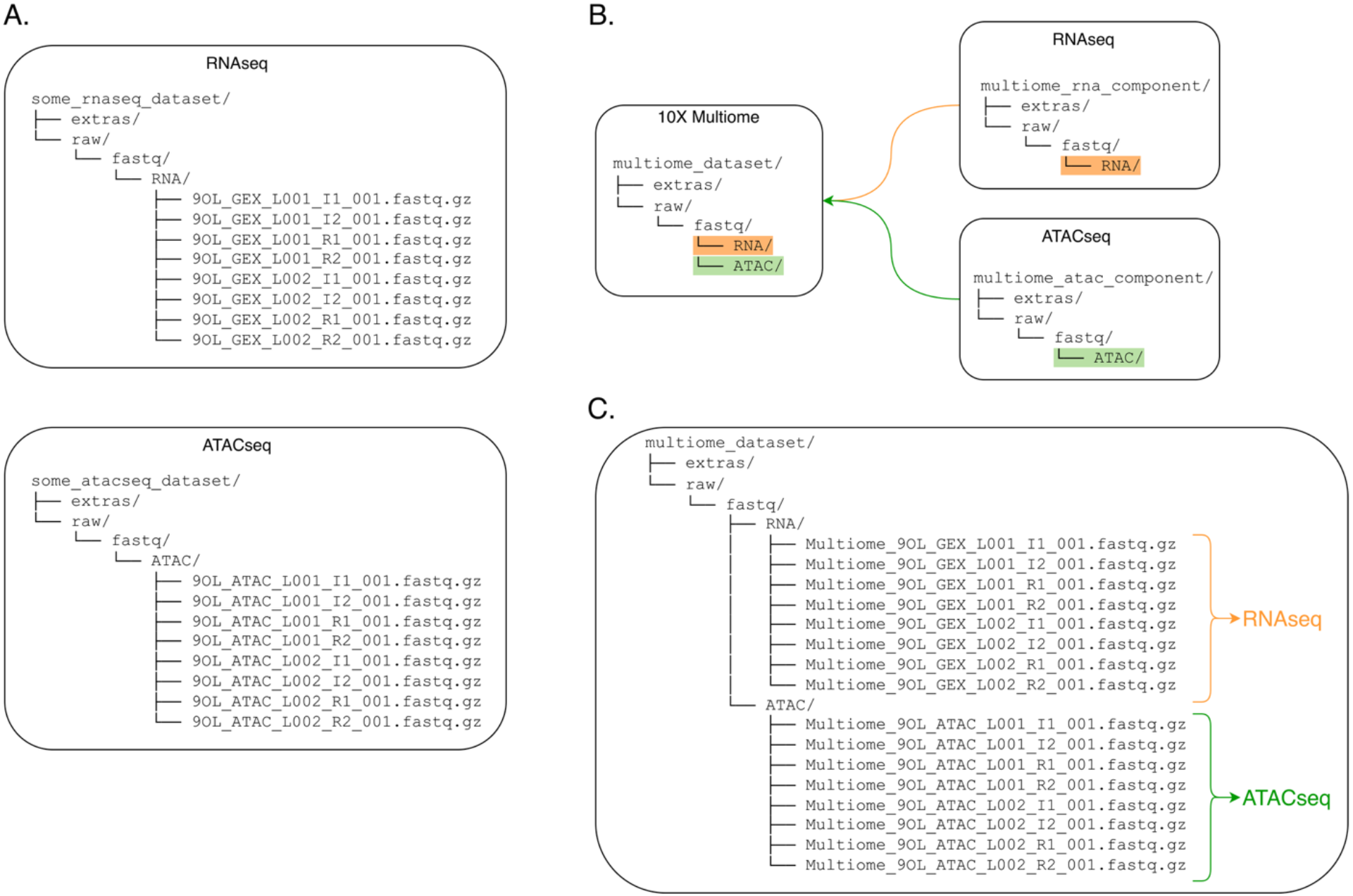
An example instance of a dataset organizational structure. The file and directory specifications provide consistent data file structures across assay types and data providers. (A) The RNA sequencing specification requires all FASTQ files to be in the “raw/fastq/RNA” directory, while the FASTQ files for DNA sequencing assays are in the “raw/fastq/ATAC” directory. (B) The 10X Genomics Multiome assay type consists of both RNA and DNA sequencing. This schematic illustrates how the RNA and DNA assay specifications are used as components in the 10X Genomics Multiome specification. (C) An example of a 10x Genomics dataset, as might be shared by a data provider.

The HuBMAP Consortium archives data for 52 different assay types (**Supplementary Table 3**), harmonized as much as is practical. For example, many 2D imaging assays predominantly include raw (unstitched) images, a single stitched composite image, and a few descriptive files (e.g., a JSON file with microscope parameters). In contrast, some newer assays can be characterized as combinations of other assays. One such complex assay is the 10x Genomics Single Cell Multiome protocol that includes both ATAC-seq and RNA-seq components. Similarly, the 10x Genomics Visium protocol includes both histology and RNA-seq components. The provenance model was designed to allow for these types of multi-tiered assays (i.e., where an assay is composed of component assays) (**Fig. 2**). We also designed the descriptive metadata schemas and file-directory structures so that assays can be pieced together. For example, a 10x Genomics Multiome dataset includes both ATAC-seq and RNA-seq components (**Fig. 4B and 4C**). In this case, the files and directories in a 10x Genomics Multiome dataset are structured such that they may be processed as a 10x Genomics Multiome dataset, an ATAC-seq dataset, or an RNA-seq dataset.

Some imaging workflows generate files that are shared across multiple datasets. For example, a single OME-TIFF^18^ image of an entire microscope slide may contain multiple tissue sections (each used for a separately-assayed dataset). The HuBMAP file organization specifications are designed to accommodate shared files in such a way as to simplify the management and uploading of data files, while still allowing for automated processing by the ingest system (see **Supplementary Figure 1**).

### 3.4 Enabling standardized outputs

The use of both standardized file structures and standardized descriptive metadata enables HuBMAP not only to ensure FAIRness of submitted datasets, but also to apply a relatively small number of standardized processing pipelines across diverse assay types. This efficient workflow, in turn, allows the Consortium to generate distinct, standards-compliant output datasets tailored to specific assay modalities and downstream analytical ecosystems. Importantly, the approach produces harmonized outputs that further enhance the findability, accessibility, interoperability, and reusability of the data.

Accordingly, HuBMAP disseminates processed data in community-standard formats appropriate to the nature of each assay type. For non-spatial transcriptomic and proteomic datasets, HuBMAP provides both raw unfiltered count data and processed expression matrices. Processed data are distributed in H5AD (AnnData)^21^ format, compatible with single-cell analysis tools such as Scanpy^22^ and Seurat^23,24^.

For spatially resolved assays^25^, HuBMAP provides OME-TIFF files for imaging-based modalities (e.g., CODEX/PhenoCycler) or SpatialData objects containing co-registered images and spatial coordinates alongside molecular measurements within a unified structure. SpatialData is a cloud-optimized, Zarr-backed format that enables scalable visualization and analysis across assay types^26^. Also provided are OME-Zarr^27^ files, used for large-scale spatial imaging data (e.g., 2.5D or 3D datasets). This cloud-optimized format divides the data into smaller chunks for faster loading and parallel processing, making it well suited for web-based interactive visualization tools, such as Vitessce^28^. All HuBMAP raw and processed datasets that are published with these standards are available on the HuBMAP Data Portal^10,29^ accessible at https://portal.hubmapconsortium.org.

### 3.5 Alignment with the Human Cell Atlas

Where possible, HuBMAP’s format choices are aligned with those adopted by complementary consortia such as the Human Cell Atlas, the Kidney Precision Medicine Project, the Cellular Senescence Project, and the Human Reference Atlas, thereby reducing barriers to cross-consortium data integration and reuse. As an example, we compare the Human Cell Atlas (HCA) data standards^30,31^ with those of HuBMAP.

The HCA is a global consortium seeking to map all cell types in the human body, and utilizes a mature and well-described metadata structure. **Figure 5** illustrates commonalities and differences between the HuBMAP and HCA provenance models^32^, using the example of a single-cell sequencing experiment. While the provenance models are comparable, the models differ around the capturing of descriptive metadata at the various stages. For example, while both consortia capture protocols for each stage in the provenance map, HuBMAP also includes extensive metadata schemas for each of the biomaterial collection steps (e.g., the collection of tissue blocks and the dissociation of tissue into suspensions). At the assay/experiment level both HCA and HuBMAP capture protocols and extensive metadata (see **Supplementary Table 6** for a comparison of RNA-seq experiment schemas). With regard to the data files, HCA focuses on the key data files (e.g., FASTQ files and cell annotations for sequencing), whereas HuBMAP seeks to capture all files pertaining to an experiment (e.g., configuration files, gene or protein panel lists) in a systematic way, enabling re-processing as improved analysis pipelines emerge. While both HCA and HuBMAP have formal specifications for capturing provenance around sequencing experiments, the HuBMAP data standards apply this same rigor to all assay types and file organization specifications.

**Figure 5.**
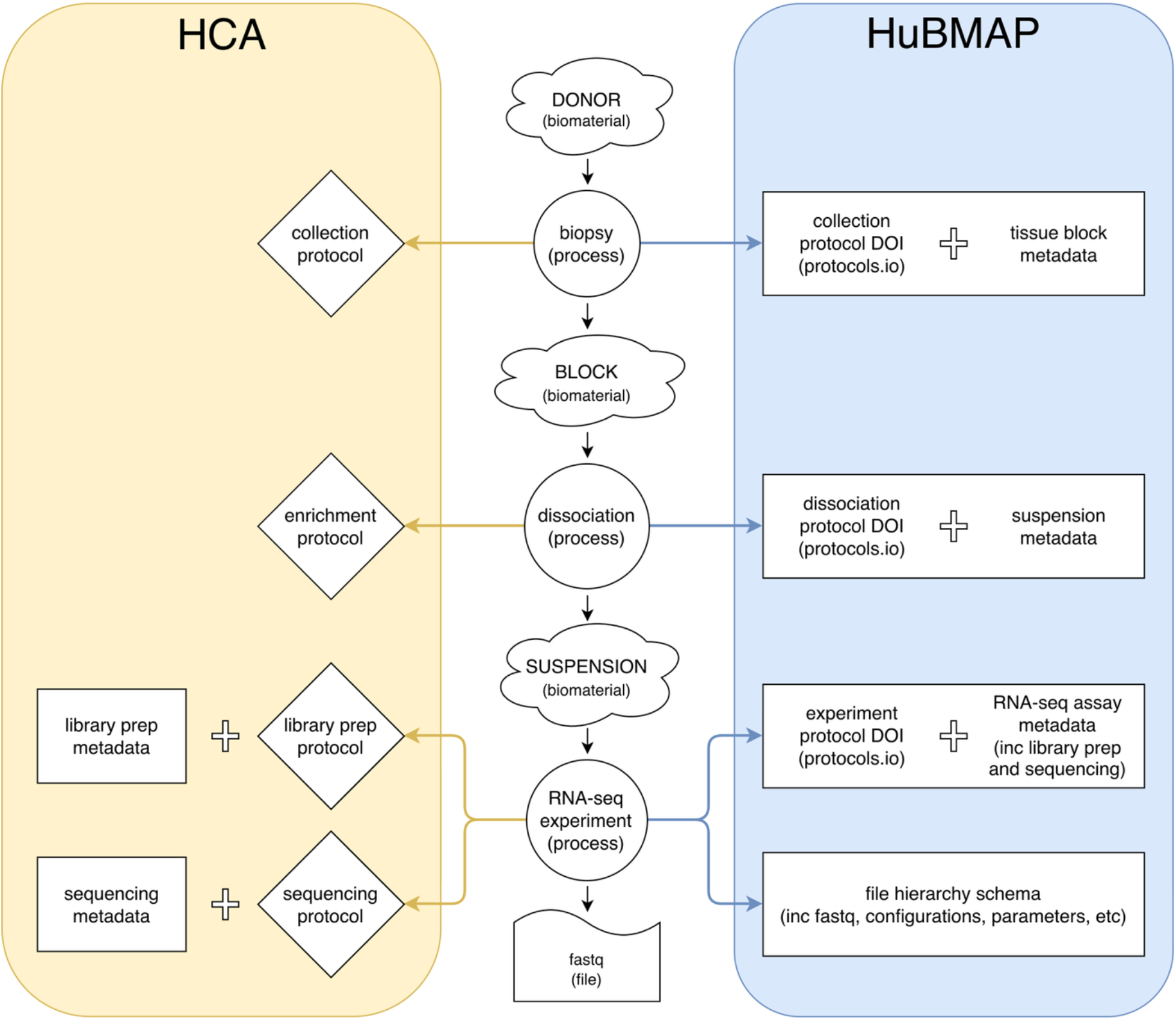
Alignment of HuBMAP and HCA provenance models. While the provenance models are closely aligned, there are major differences between the capture of metadata and data files.

### 3.6 Summary of HuBMAP Standards

The HuBMAP Consortium metadata standards include our flexible and robust data provenance model, numerous comprehensive descriptive metadata schemas, and detailed structural requirements for packaging data files for sharing. Rich metadata allow a data provider to specify all the “data about the data” needed for another investigator to understand the nature of a study and how the study was conducted. Specifying the file and directory structures enhances “accessibility” by clearly describing and enforcing the organization of the data. Together, these standards provide a formal, well-documented, human- and machine-actionable encapsulation sufficient for a third party or computer to determine what has been done and whether the data are reusable for a given context or use case.

## 4. Software Infrastructure to Support the Workflow

The HuBMAP consortium enforces adherence to its descriptive metadata standards through an information technology infrastructure that supports investigators at the point of metadata authoring and ensures metadata quality through centralized curation and validation. This infrastructure is based on the Center for Expanded Data Annotation and Retrieval (CEDAR) Workbench developed at Stanford University^33,34^. Using the CEDAR Workbench, metadata schema specifications are formalized as CEDAR templates. HuBMAP maintains a library of templates that serve as reusable blueprints defining how standards-adherent metadata should be captured for each dataset or experiment. Each CEDAR template specifies the necessary structure and semantics of metadata entries, including required fields, acceptable value sets, data types, and other validation rules, ensuring that submitted metadata are complete and consistent with the specifications. For metadata fields constrained by controlled vocabularies (e.g., ontologies or value sets), CEDAR enforces these constraints by retrieving terms from the BioPortal ontology repository^34–36^. In HuBMAP, these terms are drawn specifically from the HuBMAP Research Attributes Value Set (HRAVS, described in **Section 3.2**), which is published to BioPortal.

From a CEDAR template, the Workbench can natively generate a web form for metadata acquisition. HuBMAP, however, requires metadata to be submitted via CEDAR-generated spreadsheets. For each metadata schema, CEDAR automatically creates a corresponding Excel file that encodes the schema together with its ontological constraints^37^. These spreadsheets provide a structured interface for metadata entry, with predefined column headings corresponding to metadata attributes and, where applicable, cells constrained to controlled value sets. Each row represents a single dataset, allowing investigators to capture metadata for multiple datasets within a single file, which is convenient for experiments involving large collections of assays.

Spreadsheets offer a familiar and flexible format for investigators, lowering the barrier to metadata submission while reinforcing adherence to standards through predefined fields and controlled vocabularies. Data providers populate these spreadsheets with descriptive metadata and submit them to the data ingest system. Since manual data entry introduces the possibility of errors, automated validation is required to ensure that submitted metadata conform to the specified schemas prior to ingestion.

To help data providers comply with HuBMAP’s metadata standards, the CEDAR Spreadsheet Validator^37,38^, built on the CEDAR infrastructure, serves as a user-friendly web tool for bulk validation of metadata from one or more datasets (**Fig. 6A**). Given a metadata spreadsheet, the validator performs two key checks:

**Figure 6.**
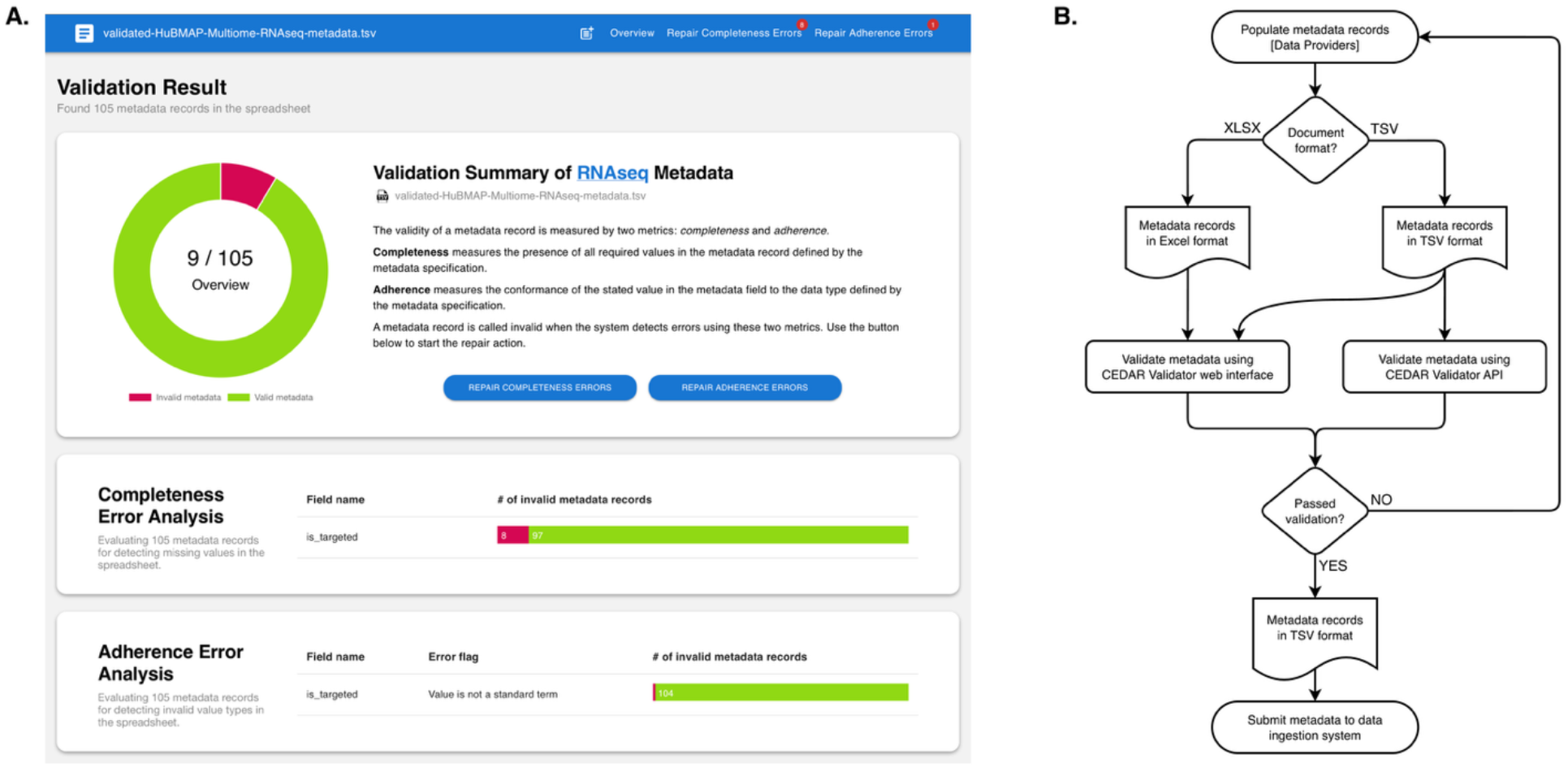
**(A)** An example screenshot from the online CEDAR metadata validation tool. After uploading a table of metadata (in Excel XLSX or tab-separated values format), the tool analyzes the supplied information and provides the user with feedback across all records and fields, with regard to completeness (i.e., were all required metadata fields included) and adherence (i.e., did each metadata value conform to its expected data type). **(B)** Schematic of metadata validation workflow. Metadata, provided as Excel (XLSX) or tab-separated values (TSV) files, are validated with the CEDAR Validator using either the web interface or through the API.

1. **Field completeness** – verifying that all required fields are filled
2. **Value adherence** – ensuring that values conform to the constraints defined in the corresponding CEDAR template, including valid entries from controlled value sets, string patterns, and correct data types.

The validator is also available through a remote API that facilitates integration with external software systems. This API enables the central HuBMAP data ingest pipeline to pre-screen metadata records before accepting dataset files for submission. Only records that pass validation proceed to subsequent checks of file organization and format. Fully validated datasets are then ingested for further refinement and enrichment. **Figure 6B** illustrates the metadata validation workflow and its integration with the data ingest system.

Together, CEDAR templates and the Metadata Validator operationalize key elements of the FAIR Guiding Principles by enforcing standardized, structured metadata at the point of submission, while remaining flexible and accessible for data providers. This combination of rigor and usability helps to ensure that metadata collected across multi-institutional research programs such as HuBMAP and SenNet are consistently structured, richly annotated, and interoperable across systems.

## 5. The Cellular Senescence Network (SenNet) use case

HuBMAP is closely aligned with a second consortium, the NIH Common Fund’s Cellular Senescence Network (SenNet) Program^39–41^, the goal of which is to comprehensively identify and characterize the differences in senescent cells across the body, across various states of human and murine health.

Given the substantial overlap of single-cell and spatial assays across the two consortia, as well as common personnel, it made sense for SenNet to adopt and adapt all three components (i.e., the provenance model, descriptive and structural metadata schemas) of HuBMAP’s existing metadata standards. In particular, the componentized design of HuBMAP’s data provenance model made it possible to accommodate SenNet’s goals despite the fact that, unlike HuBMAP, SenNet sought to collect data from mouse sources, as well as from humans. This required the development of only one new murine source metadata specification; the remaining provenance model, descriptive metadata schemas and file organization structures were immediately usable without substantive modification or new development. For example, the source provenance information for a dataset (as defined by the provenance graph) allows the downstream processing to proceed automatically, using the mouse reference genome, without changing any of the assay metadata specifications.

This adoption of the HuBMAP infrastructure allowed SenNet data providers to begin registering and uploading SenNet data in a relatively short period of time. Likewise, authorized members of both consortia benefit from the rapidly-maturing templates, tools, and documentation being co-developed across both consortia. Additionally, with both Data Coordination Working Groups maintaining a disciplined approach to the creation and maintenance of dataset reporting guidelines, new metadata specifications could be readily shared as a unified standard across consortia, regardless of which consortium motivated the change and which handled the actual development.

In the spirit of FAIR data, HuBMAP and SenNet aspire to make all their standards and tools sufficiently comprehensive to be adopted in their entirety, yet sufficiently flexible to be easily modified to meet the expanded needs of other research communities. With coordination, these data-management components create a mutual feedback loop, as is the case with SenNet and HuBMAP, who offer one another unified standards for each source, sample, and assay across both consortia—avoiding the need for each community to create its own standards from scratch.

## 6. Discussion

Since the publication of the FAIR Guiding Principles in 2016, there has been widespread recognition of the importance of making research data available for verification and reuse^1^. However, despite strong endorsement from funders, publishers, and the scientific community, many investigators remain uncertain about how to operationalize FAIRness in practice. A common misconception is that simply making data publicly available is sufficient^2^. In reality, FAIR data require more than accessibility: They depend critically on the presence of rich, standardized metadata that enable discovery, interpretation, and reuse.

Although the FAIR principles encompass 15 guidelines, many pertain to repository-level features that are not under the direct control of individual investigators. In contrast, researchers have direct responsibility for the quality of the metadata accompanying their datasets. This observation motivates the central premise of this work: that the most effective way to advance data FAIRness is to ensure that datasets are annotated with rich metadata that adhere to community standards^1,42^.

The HuBMAP Consortium operationalizes this premise through an integrated framework that combines community commitment, standards development, governance processes, and supporting infrastructure. Central to this framework is the development of comprehensive descriptive metadata and structural file organization standards that cover the diverse biological samples and experimental assays used within the consortium. These standards are established through a formal process led by the Data Coordination Working Group (DCWG), which engages relevant stakeholders and emphasizes consistency, completeness, and usability. Metadata schemas are authored in the CEDAR Workbench^13,33^, ensuring alignment with established ontologies and value sets and enabling dissemination of machine-actionable standards across the consortium. File hierarchy and naming standards are represented as regular expressions, providing a versatile, common language and allowing for automated structural validation. Using this process, HuBMAP has developed standardized data-reporting specifications for all assay types it supports, facilitating data harmonization, retrieval, and downstream analysis. In addition to enabling human understanding and reuse, such structured, standards-adherent specifications also support automated data discovery and interpretation by computational systems, including emerging AI-driven approaches that rely on consistent, machine-readable representations.

A key challenge in implementing FAIR data practices is the burden associated with generating high-quality metadata. Investigators often view metadata annotation as secondary to experimental work, and the absence of clear standards and usable tools further complicates this task. The poor quality of metadata in many public repositories reflects these challenges^3,43^. HuBMAP addresses these issues by integrating metadata capture into familiar and efficient workflows. In particular, the consortium uses the CEDAR Workbench to generate spreadsheets tailored to each assay type, with metadata fields appropriately constrained and including relevant controlled vocabularies. This approach aligns with common laboratory practices while reducing user burden and minimizing errors. Automated validation tools further ensure that submitted metadata conform to established standards. By making adherence to metadata standards straightforward and largely transparent, HuBMAP enables FAIR data practices to become a routine component of data generation rather than an additional obligation. The use of this spreadsheet-based approach has allowed HuBMAP and SenNet to collect more than 13,500 datasets from over 40 funded components (representing over 60 institutions), with confidence that the metadata adhere to standard schemas and that the data are FAIR.

The HuBMAP experience highlights primary barriers to the creation of FAIR data: (1) a lack of community-endorsed metadata standards, (2) a lack of consistent and well documented file and directory structures for data sharing, and (3) the difficulty of incorporating such standards into everyday research workflows. The consortium addresses the first two barriers through coordinated, community-driven development of metadata standards for a wide range of experimental assays. The final barrier is addressed through the deployment of user-friendly tools that embed the metadata standards directly into the data-acquisition process. Together, these efforts demonstrate that these barriers can be overcome through a combination of governance, standards, and infrastructure.

The success of this approach is reflected in its scalability, adoption, and external evaluation. HuBMAP has developed and deployed numerous data reporting standards across a large and diverse set of datasets, and investigators within the consortium have readily adopted the associated workflows. Moreover, the framework has proven transferable, as evidenced by its application in related efforts within the SenNet Consortium. Notably, in an NIH-commissioned assessment of FAIRness across 28 biomedical data repositories using more than 20 evaluation criteria, HuBMAP was the only resource to achieve a perfect score^44^. These results provide independent validation of the effectiveness of the HuBMAP approach to data management. Importantly, the HuBMAP framework is not tied to a specific scientific discipline or proprietary infrastructure. The tools used—including the CEDAR Workbench, spreadsheet-based metadata templates, and validation utilities—are open-source and broadly applicable. Research groups seeking to adopt a similar approach can begin by identifying or developing appropriate metadata and data file reporting guidelines, while using CEDAR Workbench to formalize the metadata standards. Despite increasing recognition of the importance of reporting guidelines for reproducibility and data reuse, many domains still lack such standards^1,42^.

General-purpose data repositories, such as Dryad (https://datadryad.org) and the Open Science Framework (https://osf.io), already support the use of structured metadata and can serve as practical endpoints for storing FAIR-compliant datasets^45–48^. These platforms increasingly incorporate community-developed reporting guidelines, including those originating from HuBMAP, enabling investigators to annotate their data in a standards-adherent manner at the time of submission. Thus, a complete FAIR data workflow can be implemented using existing open tools and repositories without requiring specialized infrastructure.

The HuBMAP experience demonstrates that advancing data FAIRness does not require complex or burdensome processes. Rather, it depends on community-driven standards development and practical, user-centered tools. Any laboratory or research group can make use of the existing HuBMAP framework, or may readily adapt it for their specific needs, using the resources and workflows described in this paper. By integrating these elements into routine research workflows, it is possible to make FAIR data practices both accessible and sustainable, thereby realizing the broader goals of data sharing and reuse across the scientific community.

## Data Availability

- All HuBMAP metadata reporting standards are listed in **Supplementary Table 3**.
- The HuBMAP Research Attributes Value Set (HRAVS) is available on BioPortal: https://purl.humanatlas.io/vocab/hravs.
- Datasets published using these reporting standards are available on the HuBMAP and SenNet data portals:
  - HuBMAP Data Portal: https://portal.hubmapconsortium.org/
  - SenNet Data Portal: https://data.sennetconsortium.org/

## Code Availability

All HuBMAP, SenNet, and CEDAR software used in the workflows presented in this paper is open source and freely available from https://github.com/hubmapconsortium, https://github.com/sennetconsortium, and https://github.com/metadatacenter.

## Acknowledgments

We are grateful to Ajay Pillai for providing essential vision, leadership and support in all stages of this work. This research was supported, in part, by the U.S. National Institute of Health under awards OT2OD033753, OT2OD033758, OT2OD033759, OT2OD033760, OT2OD033761, U01HD110336, U01HL166058, U54AG076041, U54AG079754, U54AR081775, U54DK134301, U54DK134302, U54EY032442, U54HD104392, U54HD104393, U54HD110347, U54HG01272302, U54HL165440, U54HL165442, U54HL165443, UG3CA256959, UG3CA275669, UG3CA275681 and UH3CA275669.

## Contributions

S.A.F., R.M. and P.M.K. led the development of the metadata and directory structure specifications. J.H., S.A.F., R.M. and B.H. contributed to terminology harmonization, specification review and refinement, and mapping to existing ontologies. M.M.R., S.Donahue, P.C. and S.G. provided software engineering and bioinformatician review of the specifications. All other authors participated as subject matter experts, in the design and review of the metadata and directory structure specifications. S.A.F. and M.A.M. led the writing of this paper, with J.H., E.N., R.M., J.C.S., P.D.B., B.H. and M.L.T. assisting with the writing. J.R., S.J., F.G., S.G., I.S.V., L.S., D.C.S., M.L.T., J.F., Y.H., P.S., M.T.E. and N.G. reviewed and commented on this paper. S.A.F., R.M. and P.M.K. provided project management and leadership during the DCWG meetings, where the specifications were drafted. S.A.F., J.C.S., P.D.B., M.A.M. and N.G. provided overall oversight and leadership. S.A.F., M.A.M. and J.C.S. are corresponding authors.

## Full list of DCWG Member authors

| First Name | Middle Name | Last Name | Affiliation |
| --- | --- | --- | --- |
| Kyung | Jin | Ahn | Children's Hospital of Philadelphia, Philadelphia, PA, USA |
| Christopher | R | Anderton | Pacific Northwest National Laboratory, Richland, WA, USA |
| Shovik |  | Bandyopadhyay | Children's Hospital of Philadelphia, Philadelphia, PA, USA;<br>University of Pennsylvania, Philadelphia, PA, USA |
| Kenneth | C | Bedi | University of Pennsylvania, Philadelphia PA, USA |
| Maigan |  | Brusko | University of Florida, Gainesville, FL, USA |
| Martha |  | Campbell-Thompson | University of Florida, Gainesville, FL, USA |
| James |  | Carson | The University of Texas at Austin, Austin, TX, USA |
| Chase | M | Carver | Mayo Clinic, Rochester, MN, USA |
| Jing |  | Chen | University of Florida, Gainesville, FL, USA |
| Anthony | M | Corbett | University of Rochester Medical Center, Rochester, NY, USA |
| Alexandra | E | Cuaycal | University of Florida, Gainesville, FL, USA |
| Penny |  | Cuda | Carnegie Mellon University, Pittsburgh, PA, USA |
| Dinh |  | Diep | University of California San Diego, La Jolla, CA, USA<br>Currently: Altos Labs, San Diego, CA, USA |
| Sergii |  | Domanskyi | The Jackson Laboratory for Genomic Medicine, Farmington, CT, USA |
| Sean |  | Donahue | Carnegie Mellon University, Pittsburgh, PA, USA |
| Michael |  | Duffy | University of Pennsylvania, Philadelphia PA, USA |
| Michael | T | Eadon | Indiana University School of Medicine, Indianapolis, IN, USA |
| Jean |  | Fan | Johns Hopkins University, Baltimore, MD, USA |
| Melissa | A | Farrow | Vanderbilt University, Nashville, TN, USA |
| Kathleen | M | Fisch | University of California San Diego, La Jolla, CA, USA |
| William | F | Flynn | The Jackson Laboratory for Genomic Medicine, Farmington, CT, USA |
| James | M | Fulcher | Pacific Northwest National Laboratory, Richland, WA, USA |
| Nils |  | Gehlenborg | Harvard Medical School, Boston, MA, USA |
| Soumya |  | Ghose | GE HealthCare, Niskayuna, NY, USA |
| Fiona |  | Ginty | GE HealthCare, Niskayuna, NY, USA |
| Joana | P | Gonçalves | Delft University of Technology, Delft, The Netherlands |
| Yongqun |  | He | University of Michigan, Ann Arbor, MI, USA |
| Po |  | Hu | Children's Hospital of Philadelphia, Philadelphia, PA, USA;<br>University of Pennsylvania, Philadelphia, PA, USA |
| Sanjay |  | Jain | Washington University School of Medicine, St. Louis, MO, USA |
| Thomas | V | Karathanos | Stanford University, Stanford, CA, USA |
| Madhurima |  | Kaushal | Washington University School of Medicine |
| Angela | RS | Kruse | Vanderbilt University, Nashville, TN, USA<br>Currently: The Ohio State University, Columbus, OH, USA |
| Yumi |  | Kwon | Pacific Northwest National Laboratory, Richland, WA, USA |
| Blue | B | Lake | University of California San Diego, La Jolla, CA, USA<br>Currently: Altos Labs, San Diego, CA, USA |
| Roy |  | Lardenoije | Delft University of Technology, Delft, The Netherlands |
| Shin |  | Lin | Emory University, Atlanta, GA, USA |
| Yiing |  | Lin | Washington University, St. Louis, MO, USA |
| Scott | A | Lindsay | University of California, San Diego, CA, USA |
| Peiran |  | Lu | Children's Hospital of Philadelphia, Philadelphia, PA, USA;<br>University of Pennsylvania, Philadelphia, PA, USA |
| Clayton |  | Mathews | University of Florida, Gainesville, FL, USA |
| Elizabeth |  | McDonough | GE HealthCare, Niskayuna, NY, USA |
| Ricardo |  | Melo Ferreira | Indiana University School of Medicine, Indianapolis, IN, USA |
| Emma | M | Monte | Stanford University, Stanford, CA, USA |
| Kathleen |  | O'Neill | University of Pennsylvania, Philadelphia PA, USA |
| Minxing |  | Pang | University of Pennsylvania, Philadelphia PA, USA |
| Mana |  | Parast | University of California San Diego, La Jolla, CA, USA |
| Liming |  | Pei | Children's Hospital of Philadelphia, Philadelphia, PA, USA;<br>University of Pennsylvania, Philadelphia, PA, USA |
| Samuel |  | Peters | University of Minnesota, Minneapolis, MN, USA |
| Gloria |  | Pryhuber | University of Rochester Medical Center, Rochester, NY, USA |
| Ling |  | Qin | University of Pennsylvania, Philadelphia PA, USA |
| Presha |  | Rajbhandari | Columbia University, NYC, NY, USA |
| Matthew | M | Ruffalo | Carnegie Mellon University, Pittsburgh, PA, USA |
| Pinaki |  | Sarder | University of Florida, Gainesville, FL, USA |
| Diane | C | Saunders | Ann & Robert H. Lurie Children's Hospital of Chicago, Chicago, IL, USA; Northwestern University, Chicago, IL, USA |
| Kevin |  | Schneider | Buck Institute, Novato, CA, USA |
| Lingyan |  | Shi | University of California San Diego, La Jolla, CA, USA |
| Santhosh |  | Sivajothi | The Jackson Laboratory for Genomic Medicine, Farmington, CT, USA |
| David |  | Smith | Children's Hospital of Philadelphia, Philadelphia, PA, USA |
| Jeff | M | Spraggins | Vanderbilt University, Nashville, TN, USA |
| Valentina |  | Stanley | University of California, San Diego, CA, USA |
| Kai |  | Tan | Children's Hospital of Philadelphia, Philadelphia, PA, USA; University of Pennsylvania, Philadelphia, PA, USA |
| Anusha |  | Thadi | Children's Hospital of Philadelphia, Philadelphia, PA, USA |
| Hua |  | Tian | Columbia University Medical Center, NYC, NY, USA |
| Morgan | L | Turner | Harvard Medical School, Boston, MA, USA |
| Ioannis | S | Vlachos | Beth Israel Deaconess Medical Center, Boston, MA, USA; Harvard Cancer Center, Boston, MA, USA; Broad Institute of MIT and Harvard, Boston, MA, USA |
| Seth |  | Winfree | University of Nebraska Medical Center, Omaha, NE, USA<br>Currently: QCDx Inc, Farmington, CT, USA |
| Pei-Hsun |  | Wu | Johns Hopkins University, Baltimore, MD, USA |
| Kevin | J | Zemaitis | Pacific Northwest National Laboratory, Richland, WA, USA |
| Mowei |  | Zhou | Pacific Northwest National Laboratory, Richland, WA, USA<br>Currently: Zhejiang University, Hangzhou, Zhejiang 310058, China |
| Chenchen |  | Zhu | Stanford University, Stanford, CA, USA |

