## Supplemental Information for "The HuBMAP Framework for Advancing Data FAIRness"

### Supplementary Information

#### Supplementary Description 1. Biological samples consist of tissue blocks, sections, and suspensions.

**Blocks** are pieces of tissue typically sized to fit into tissue cassettes or freezer molds, appropriate for storage or sectioning, although a block could be used to represent an entire organ. The blocking of tissue is usually done by hand with a scalpel, resulting in a thick piece of tissue (typical Z plane 0.5-1 cm thick). Organs, “organ pieces,” and biopsies are considered blocks. **Sections** are cut from blocks to achieve much thinner samples usually placed on slides, in cultures, or pulverized for isolation of nuclei. Sections are typically 1-60 µm thick but can be up to about 1 mm for 3D imaging or short-term cultures. Sections generally represent a final assayed sample, not cut any thinner before use (for imaging, culture, extraction or dissociation). **Suspensions** are dissociated cells, nuclei or cellular membrane-bound organelles in a liquid suspension.

#### Supplementary Description 2. Examples of how to capture the temporal relationship between entities.

Given the complexity of capturing temporal durations, specific examples are offered to provide additional clarity. For example, at one HuBMAP research site, knee blocks are received from a tissue bank on wet ice within 48 hours after death. “Warm ischemic time” is the time from when death was reported to when the tissue bank received the tissue, whereas “cold ischemic time” is the time from when the tissue bank received the tissue to when the knee blocks are received in the lab. The “source storage time” for the suspension is how long the knee block remained on ice, in the laboratory, before being opened for cartilage harvesting. The “processing time” for the suspension is the length of time from when a joint capsule is opened to when the digested cartilage is placed into a buffer solution. At another HuBMAP site, a donor bank provides the entire female reproductive system in the operating room, which is then transported to a research laboratory. The warm ischemic time, which measures the duration between time of death and the start of cold profusion, is provided by the donor bank. Cold ischemic time begins upon starting cold profusion and ends when the HuBMAP team gets the organs back to their laboratory. Once in the laboratory, the organs are immediately dissected into tissue blocks. In this case, the “source storage time” for the tissue blocks is zero, since the blocks are derived immediately from the organ source. The “processing time” for tissue blocks is the length of time from when the organ is received to when the tissue block is either embedded in OCT (optimal cutting temperature) compound, or snap-frozen and placed in a freezer for later processing.

#### Supplementary Table 1. Dataset Subtyping Rules

For some datasets, the value of the *dataset_type* metadata field is not sufficient to uniquely identify the dataset class. For example, there are many types of datasets that have the “RNAseq” *dataset_type* value, and some of these have different downstream analysis requirements. In these cases, the values of specific additional metadata fields are used to uniquely determine the dataset subtype. The Dataset Subtyping Rules are also used to validate metadata values for specific dataset types (e.g., the *barcode_size* must be consistent with the *oligo_probe_panel*). The specific metadata fields and values used by HuBMAP Consortium are included in this table. Note that grayed-out cells in the table denote fields that are not relevant to the workflow decision for a given assay, while “Not applicable” is a valid metadata value for certain fields, which may be relevant when choosing a workflow.

#### Supplementary Table 2. Harmonizing across sequencing assays.

The various sequencing assays proved to be the most complicated assays to characterize. Our goal for the RNA-seq metadata schema was to be general enough for the schema to work for all major variants of next-generation RNA sequencing assays, but specific enough to delineate among RNA-seq variants for downstream processing of primary data. To achieve our goal, we first created separate schemas for a few variants of RNA-seq assays (e.g., SmartSeq and 10x Genomics technologies) with different input entities (e.g., bulk vs. single cells). This allowed us to better identify the critical components for each variant. We then collapsed these schemas into two RNA-seq schemas, based on whether or not the assay used probe panels (i.e., oligonucleotide sequences). Due to the variability in RNA-seq assay methodologies, while some metadata fields are critical to capture for a subset of technologies, these same fields are not relevant to other technologies. We thus opted to make any such fields required and categorical, allowing for “not applicable” as an option. This circumvents a situation where an investigator might miss filling in an optional field, failing to note the significance of the field. This table details the fields for the schemas that include RNA and DNA sequencing, illustrating how we harmonized across the sequencing types, with comprehensive schemas.

#### Supplementary Table 3. Data Reporting Standards

This table provides a comprehensive inventory of the data reporting standards currently defined within the HuBMAP framework. The standards are organized by category, and each includes a persistent URL along with a brief description of scope and purpose. Persistent URLs provide details for each standard, including a metadata schema and, where relevant, a file hierarchy schema. The majority of standards correspond to assay-specific metadata models, reflecting the breadth of technologies supported within the HuBMAP provenance framework. Metadata schemas for the various biosample types (block, section, and suspension) are also included. Finally, there are metadata schemas for two auxiliary files, used to describe: 1) the antibodies used in a proteomics assay, and 2) the list of individuals who contributed work toward executing the assay and generating the data. Together, these reporting standards encompass the hierarchical HuBMAP metadata and file hierarchy strategy, providing human- and machine-accessible documentation.

#### Supplementary Table 4. We created minimum specifications for the XML header of all OME-TIFF files.

Another important component of the HuBMAP Consortium reporting standards is specifications for image files. Specifically, HuBMAP adopted OME-TIFF files as the “standard” imaging assay deliverable. In this way, we seek to harmonize across imaging assays. Further, we adopted the Open Microscope Environment (OME) data model and XML file. Specifically, we built on their OME-XML minimum specification with regard to the required fields in the OME-TIFF XML file header. We chose this model to ensure that each OME-TIFF data file included the metadata required for displaying and interpreting the image. The following table denotes the fields from the OME data model that we require.

#### Supplementary Table 5. A “channels” file is typically included with OME-TIFF images, to provide additional details about the channels in the image.

OME-TIFF image files can include one or more images (a.k.a. channels). In spatial assays, each fluorophore is typically captured as a separate image and hence encoded in a separate channel. A standardized comma-separated-values (CSV) file is used to provide details about each of the channels, to facilitate downstream analyses. Each field in the CSV file is detailed in this table.

#### Supplementary Table 6. Comparing Human Cell Atlas and HuBMAP assay-level metadata schemas

This table presents a detailed, field-level comparison of sequencing assay metadata schemas defined by the Human Cell Atlas (HCA) and HuBMAP, with a focus on single-cell RNA sequencing as a representative use case. Metadata fields are grouped by conceptual categories — such as schema identifiers, dataset identification, instrument and library information, barcode/UMI details, and provenance — to show where the two approaches align and diverge. While HCA and HuBMAP offer comparable depth for RNA-seq assay metadata in many categories, the table highlights differences in how each consortium structures and names fields, the granularity of captured parameters (e.g., library QC metrics or reagent details), and the presence of pre-analytical and design variables in the HuBMAP schema.

#### Supplementary Figure 1. Ingesting data when multiple tissue sections are on the same imaging slide.

In many imaging assays, it can be advantageous to capture multiple datasets in a single imaging run. For example, multiple tissue sections can be mounted on the same imaging slide, such that all of the sections can be imaged together, often saving investigators both time and money. Some assays, like NanoString’s GeoMx, are designed to capture a hundred or more datasets in a single run. In order to accommodate these technologies and techniques, the HuBMAP reporting standards include a mechanism to simplify the organizational burden on data providers while allowing for automated validation and handling of the resulting, possibly independent, datasets.

For example, suppose a data provider mounted three tissue sections onto one microscope slide and then imaged the slide. Each tissue section was used for a different dataset, but all of the datasets share the same (whole-slide) raw and processed image files. For such cases, all of the data for the slide is grouped into a particular, standardized file hierarchy and uploaded together, to be ingested as a unit. HuBMAP requires the data provider to include a set of small GeoJSON files (https://geojson.org/), each of which delineates the region of the slide corresponding to one of the tissue sections. The raw and processed image files are placed within a “global” directory tree, since they pertain to the entire slide and thus are shared across all of the datasets. Conversely, all of the dataset-specific GeoJSON files are included in a “non_global” directory tree, since each of these files relates to a single dataset. In these shared-data uploads, a “non_global_files” metadata field is used to determine which of the “non_global” files are to be included in each dataset. During the HuBMAP data ingest process, the three datasets are automatically redistributed into separate instances of the appropriate single-dataset file hierarchy structure, for downstream use.

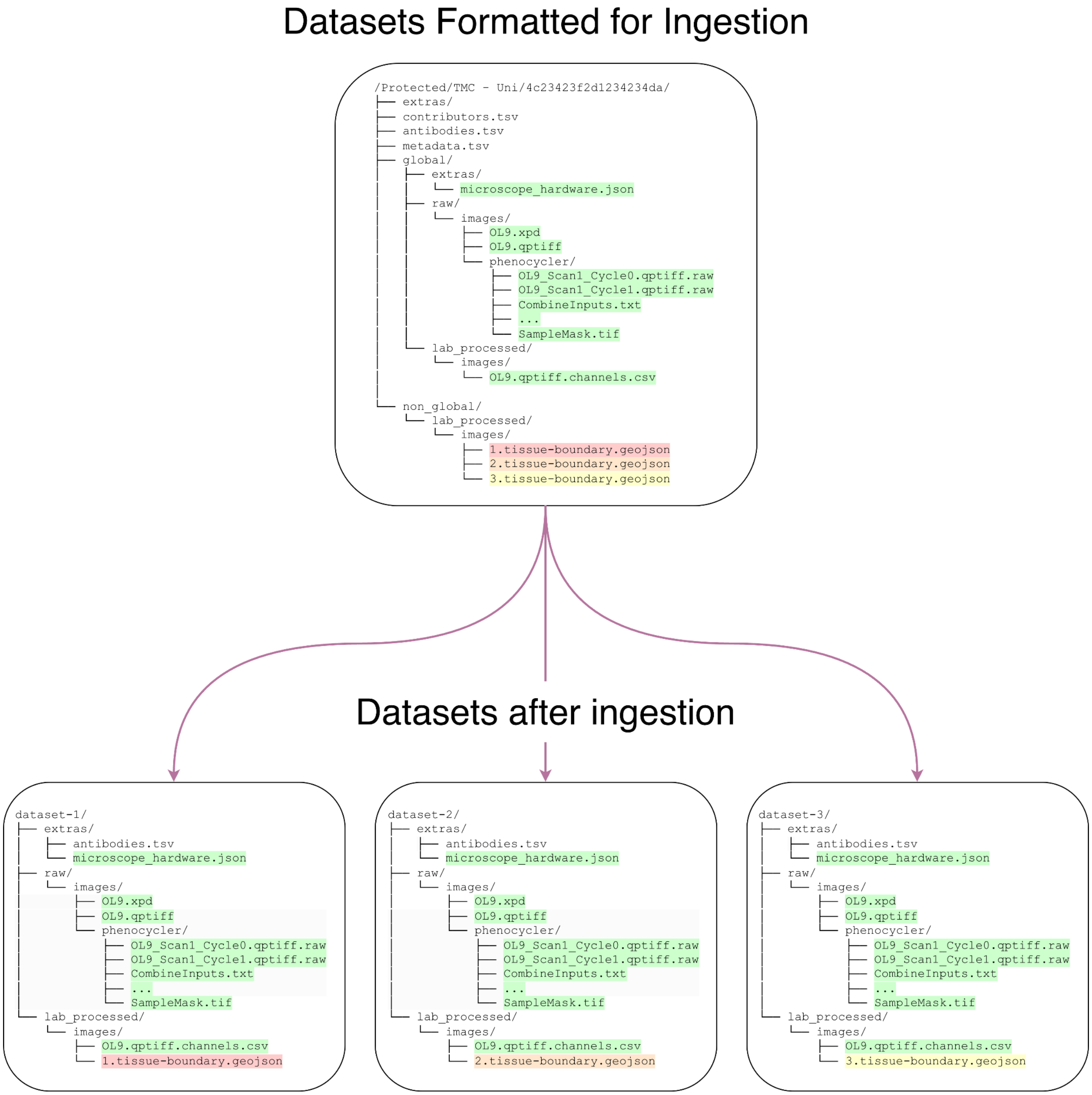

In this example, the top panel illustrates the file directory tree that would be uploaded by a data provider, in the case of an imaging experiment where three unique tissue sections were imaged together on the same slide. The “global” directory contains the files shared across each of the three datasets (i.e., the whole-slide image files and imaging parameter descriptions). The “non_global” directory contains the files that are unique to each of the final datasets. After the combined upload package is ingested by HuBMAP, it is split into the three separate datasets (lower three panels).
